# A comprehensive mycofloral diversity of pedosphere, phyllosphere and aerosphere of Som. (*Persea bombycina* Kost.)

**DOI:** 10.1101/349167

**Authors:** Manjit Kumar Ray, Piyush Kumar Mishra, Saurav Das

**Affiliations:** DBT-Advanced Institutional Biotech Hub, Bholanath College, Dhubri, Assam, India.

**Keywords:** Sericulture, Muga Silk, Diversity, Som, Mycoflora, Seasonal

## Abstract

Sericulture is an important cottage industry of Assam producing natural silk of both Mulberry and Non-Mulberry silk. The sericulture industry is closely associated with the Assamese traditions and rituals. In non-mulberry silk, Muga silkworm which is endemic to North-East India only produce exquisite silks of golden color. Rearing of Muga silkworm is one of the important aspects to producing silk of high quality. The quality of the primary host plant i.e. Som (*Persea bombycina* Kost.) greatly affects the quality of cocoon and silk produced by these industries. Therefore, planting and growing a disease-free plant has its own importance in sericulture. However, Som is very susceptible to different foliar diseases caused by fungi, which can reduce the yield of leaf from 13.8 - 41.6% annually. Therefore, a comprehensive mycofloral study of the host plant is important to forecast the future diseases and design different disease management procedures. This study has been done for a period of two years from 2014 - 2016 in Goalpara district of Assam and overall mycoflora of pedosphere (rhizosphere and non-rhizosphere), phyllosphere and aerosphere were identified and correlated with the seasonal variation. The rhizosphere, air, and phylloplane was dominated by *Rhizopus stolonifer* (22.13%; 15.08; 24.01) while *Aspergillus niger* (12.63%) was the dominant flora of non-rhizosphere. Seasonal variation was found to play an important role in shaping the mycofloral community structure in soil and phyllosphere. In summer, soil was majorly dominated by *Aspergillus niger*, *Aspergillus fumigatus*, and *Curvularia lunata* while *Rhizopus stolonifer*, *Aspergillus clavatus*, *Penicillium chrysogenum* dominated the winter soil. In phyllosphere, the Chatuwa and Jaruwa, the winter generations were mainly dominated by *Rhizopus stolonifer*. While as the environmental temperature gradually increased with relative humidity in Aheruwa, Bhodia, Kotia and Jethuwa generation there was a shift in diversity with a gradual increase in occurance of *Alternaria alternata*, *Aspergillus niger*, and *Aspergillus flavus. Pestaloptiopsis disseminata* one of the major pathogen of Som was found highest in aerosphere followed by phyllosphere and it was only dominated in Aheruwa generation. It was found that occurrence of *P. disseminata* were high when the temperature ranges between 25° - 28° with 70 −80% of relative humidity. This study provides a deep insight into the fungal diversity of host plant Som with respect to pedosphere, aerosphere, and phyllosphere and this knowledge can be used to better select the plantation area and design different disease management strategies to sustain and proliferate the industry for socio-economic development and to conserve its cultural essence.

## 1.0 Introduction

Som, the primary foodplants of Muga silkworm is a medium-sized evergreen tree which belongs to the family *Lauraceae*. It is distributed across Brahmaputra valley up to an elevation of about 500 meters. It is scantily distributed across Assam and extended to Khasi and Jayantia Hills, Meghalaya and along the lower Himalaya, it is distributed as far as up to West of Nepal (Kanjilal et al., 1992; Rahman et al., 2012). The nutritive value of the Som leaf has a considerable influence on the growth and development of silkworms. (Khanikar and Unni, 2006) reported that better the quality of the leaves of the host plant, greater the possibility of obtaining good quality cocoons. Similarly, (Chakravorty et al., 2006) mentioned, the growth of silkworm, cocoon quality and quantity of raw silk entirely depends upon the quality of leaves. Diseases, unfavorable weather conditions, insect pests, poor agronomical practices, unwanted weeds are the main reasons for low productivity. Som is vulnerable to many foliar diseases that affect the normal growth of the plant, quantity, and quality of leaves and ultimately cocoon production (Das et al., 2003)

The number of sericulture village in North-East region is about 38,000 and approximately 1.9 lakh families are engaged in this industry in Assam. According to (Unni et al., 2009), Assam is the only state in the country producing all the varieties of silk. Muga silk culture is practiced in the districts of upper Assam and certain parts of lower Assam. In lower Assam, the Goalpara district produces 800 lakhs of Muga cocoon (Baruah, 2017). Goalpara district is situated on the south bank of river Brahmaputra, and it covers an area of 1,824 km^2^ which is bounded by West and East Garo Hills districts of Meghalaya on the South, Kamrup district on the East, Dhubri district on the West and, River Brahmaputra all along the North. The geographical location of the district is between latitudes 25.53° to 26.30° North and longitudes 90.07° to 91.05° East. The agro-climatic conditions of the district are suitable for various agricultural activities. Sericulture in Goalpara district existed as a practice amongst people for long. Around 290 villages in the district are involved in sericulture activities (Kanjilal et al., 1992).(Goswami and Bhattacharya, 2013) mentioned that due to the climate suitable for silkworm rearing Goalpara district has been given the geographical identification mark. According to the report of Regional Muga Research Station (RMRS, 2000), the Eastern and Western part of Goalpara district and Southwest part of Kamrup district of lower Assam constitute important Muga growing areas where mostly seed cocoons are produced. But in recent time, rapid urbanization and climate change have affected the Som plantation very much. Migration of new pest and pathogenic dissemination has reduced the quality of leaves and as well as affected sericulture around the district. Diseases like Grey blight (pathogen –*Pestalotiopsis disseminata*) (Bharali, 1969), Anthracnose (pathogen –*Colletotrichum gloeosporioides*) (Das et al., 2005), Leaf spot (pathogen –*Phyllosticta perseae*) (Das et al., 2003), and red rust (pathogen –*Cephaleuros parasiticus*) (Thangavelu et al., 1998) are the limiting factors for the yield of Som in the particular area. So, a comprehensive study of fungal distribution is necessary to take up a future preventive measure to sustain the industry in the particular area. In this paper, we have reported the fungal distribution in the rhizospheric, non-rhizospheric soil, air and phylloplane of the plantation area of the district and determined how different environmental factors and soil parameter can affect the fungal population.

## 2.0 Materials and Methods

### 2.1 Sample collection

#### 2.1.1 Study site and duration of the study

The study was conducted at the Muga food plantation area of the Goalpara district of Assam. Five major Muga silkworm rearing villages of Goalpara district were selected *viz*. Dorapara Agia (26°5’31.525′N, 90°33’57.109′E), Budlung pahar (25°59’48.254′N,90°57’15.076′E), Lengopara (26°6’7.699′N, 90°47’9.044′E), Buraburi (25°58’ 47.057′N, 90°47’32.217′E) and Bhalukdubi Kalyanpur (26°5’38.357′N, 90°33’43.882′E) respectively (**Fig. 1**). The present study was conducted for a period of two years from February, 2014 - January, 2016.

#### 2.1.2 Soil Sample

For the collection of a rhizospheric soil sample, each Som plantlet was carefully uprooted and the soil adhering to the roots was gently shaken into a sterile polybag; the bag was tied and labeled. The non-rhizospheric soil samples were collected by digging 25 cm deep dug into the field with a sterile hand trowel; the soil was collected in sterile polybags and carefully labeled. The collected soil samples, both rhizospheric and non-rhizospheric soils were divided into four different groups based on the age of the plantlet from the surrounding of which the samples were collected *viz*. S1: 0 - 3 months; S2: 3 - 6 months; S3: 6 - 9 months and S4: 9 - 12 months.

#### 2.1.3 Air Sample

Air samples were collected using the method described by (Aneja, 2012) with petri plates containing Potato Dextrose Agar (PDA), Martins Rose Bengal Agar (MRBA) and Czapek’s Dox Agar medium supplemented with Chloramphenicol (250 mg/ml) to prevent bacterial growth. Air samples were collected for the duration of two years in six Muga crop seasons, from February 2014 to January 2016. The six different generation of Muga silkworm were as, Chatuwa (February - March), Jethuwa (April - May), Aheruwa (June - July), Bhodia (August - September), Kotia (October - November) and Jaruwa (December - January).

#### 2.1.4 Leaf Sample

Leaves of different age groups *viz*. tender, semi-mature and mature were randomly collected during rearing (outdoor) season of Muga silkworm in sterile polybags.

**Fig. 1:**
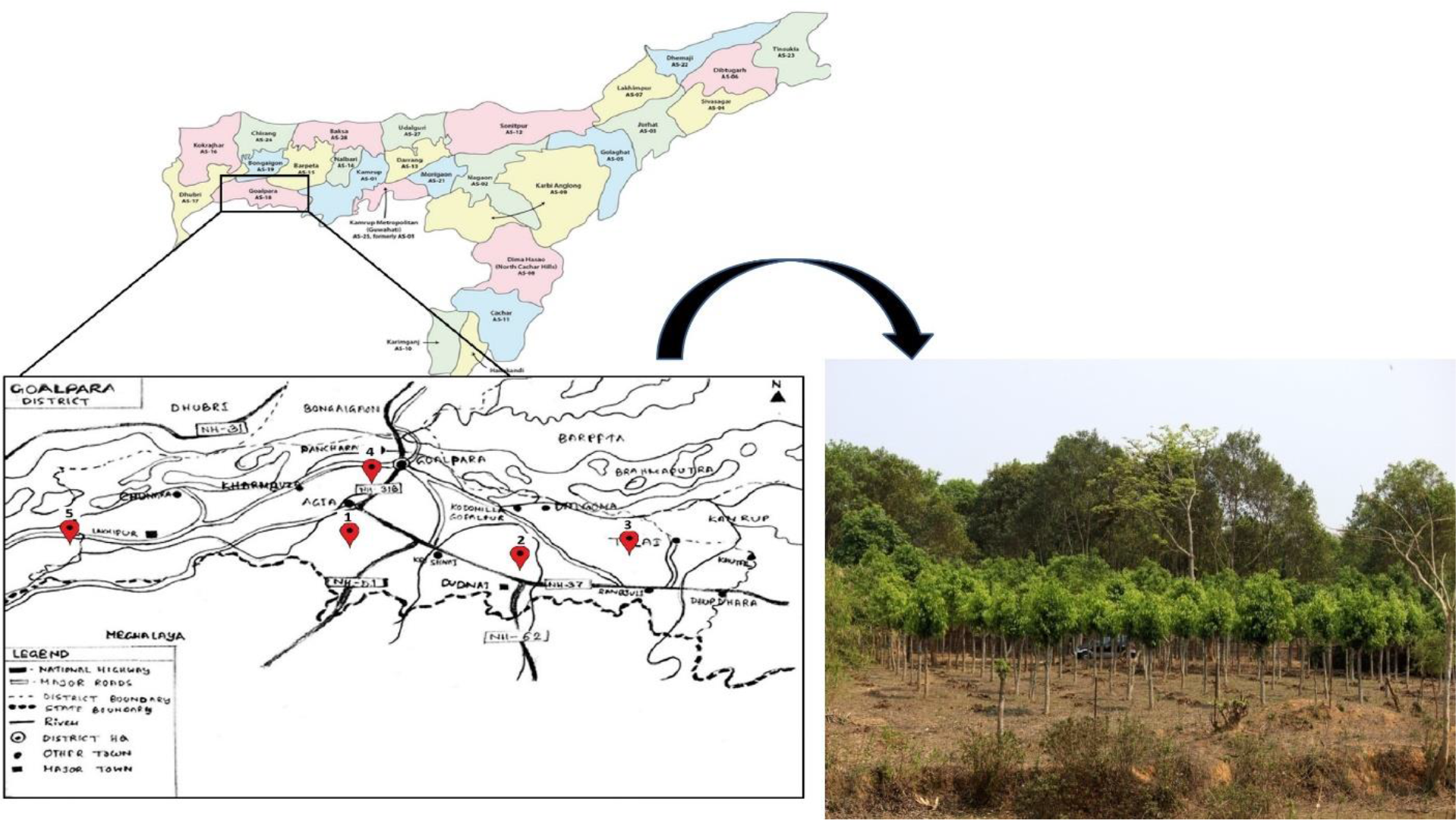
Map of the study sites with plantation spot. The red dots denote the selected sites accordingly, 1. Dorapara Agia, 2. Budlung pahar, 3. Lengopara, 4. Buraburi and 5. Bhalukdubi Kalyanpur.

### 2.2 Isolation of fungi

- Isolation of fungi from rhizospheric and non-rhizospheric soil was done by agar plating method (Atlas and Parks, 1997). The Petri dishes were incubated at 28±2° C for 7 days and then the plates were examined for the development of fungal colonies.
- The plates settled with air samples were kept for incubation at 28° ±2° C for a period of 7 days and then the plates were examined for the development of fungal colonies.
- The collected leaves were washed using serial washing technique as described by (Williams et al., 1965). Leaf discs of 1mm size were cut from each leaf with the help of sterilized borer and the dorsal, ventral surface of the cutout leaf discs was impregnated on the PDA media and incubated at 28° ± 2° C for 7 days.

### 2.3 Identification of fungi

The mycelia and spore characters of fungi were studied under microscope (Labomed, Germany) using Lactophenol cotton blue staining and identification of fungi were done with the help of “A manual of soil fungi”, by Gilman, (1959) and “Illustrated genera of imperfect fungi” by Barnett and Hunter, (1998).

### 2.4 Statistical analysis

The percentage of occurrence or contribution of each fungi to the diversity was calculated by,

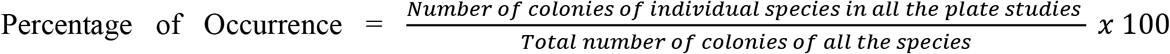

(Aneja, 2012). The compositional difference in mycofloral of rhizospheric, non-rhizospheric soil sample, air and phylloplane were represented by heatmap constructed with R software (ver. 3.2; package: vegan). The common taxa among the four environments were represented by Venn diagram drawn with software Venny (Oliveros, 2015). Diversity index (Simpson diversity index) was calculated with PAST software (ver. 3.0) to determine the richness in each environment. Canonical correspondence analysis was done using PAST software to determine the effect of environmental factors on the occurrence and diversity of mycoflora (Ryan et al., 2001).

## 3.0 Results

### 3.1 Fungal Diversity

#### 3.1.1 Soil Mycoflora

A significant difference was observed among the fungal diversity of rhizospheric and non-rhizospheric soil of Som plant. Rhizospheric soil (Simpson index - 0.9277) was rich in diversity in comparison to the non-rhizospheric samples (Simpson index - 0.8735) (**Supplementary Fig. S1**). The major dominant fungal species identified in the rhizospheric samples were, *Rhizopus stolonifer* (22.13%), *Aspergillus niger* (16.13%), *Aspergillus flavus* (12.12%), *Aspergillus fumigatus* (11.12%), *Penicillium chrysogenum* (10.88%) (**Fig. 2b**). The dominant species of the non-rhizospheric samples were *Aspergillus niger* (12.63%), *Aspergillus fumigatus (11.25%), Rhizopus stolonifer* (11.12%), *Penicillium chrysogenum* (9.5%), *Aspergillus flavus* (9.25%) (**Fig. 2a**).

Both the rhizospheric and non-rhizospheric soil samples were pre-divided into four different groups based on the age of the Som plantlets. The non-rhizospheric soil samples collected from three month age group (S1) had dominance of *Rhizopus stolonifer*, *Aspergillus niger*, *Penicillium chrysogenum*, *Mucor hiemalis*, *Trichoderma viridae* and *Saccharomyces cerevisiae;* 6 months age group (S2) had dominant mycoflora *viz. R. stolonifer*, *A. niger*, *A. fumigatus*, *T. viridae* and *P. chrysogenum*. Likewise, the non-rhizospheric sample collected from the surroundings of 9 months age group plants (S3) showed the dominance of *A. fumigatus*, *R. stolonifer*, *P. chrysogenum*, *S. cereviseae*, *T. viridae*, *A. niger*, *M. hiemalis*, *P. disseminata*, *Mycelia sterila (white)*, *C. cladosporioides and A. flavus* while in 12 months age group of plants (S4) *A. fumigatus*, *R. stolonifer*, *P. chrysogenum*, *C. cladosporioides*, *A. flavus*, *A. niger* were the dominant mycoflora (**Fig. 2a**). In rhizosphere of 3 months age group of plants (S1) dominant mycoflora were *P. chrysogenum*, *R. stolonifer*, *A. niger*, *T. viridae*, *S. cerevisae*, *M. hiemalis*, *F. oxysporum*and *M. sterila*(white), in 6 months age group of plants (S2) the dominant mycoflora were *R. stolonifer*, *P. chrysogenum*, *A. niger*, *T. viridae*, *S. cerevisae and A. fumigatus*, in 9 months old plantlet (S3) the dominant mycoflora were *R stolonifer*, *A. flavus*, *A. niger*, *C. lunata*, *P. chrysogenum*, *A. fumigatus*, *A. alternata*, *M. sterila* (white) and *T. viridae*. In 12 months age group of plants (S4) *R. stolonifer*, *A. flavus*, *A. niger*, *C. lunata*, *M. hiemalis*, *P. chrysogenum*, *A. fumigatus*, *C. cladosporioides*, *M. sterila* (white), *T. viridae* and *Trichothecium sp*. were the dominant rhizosphere mycoflora (**Fig. 2b**).

#### 3.1.2 Air mycoflora

The major aeromycoflora were *Aspergillus niger*, *Cladosporium cladosporioides*, *Rhizopus stolonifer* and *Fusarium oxysporum* during Chatuwa generation (February - March); *Rhizopus stolonifer*, *Alternaria alternata*, *Fusarium oxysporum* and *Curvularia lunata* during Jethuwa generation (April - May); *Aspergillus flavus*, *R. stolonifer* and *C. lunata* during Aheruwa (June - July); *A. flavus*, *A. fumigatus*, *A. niger*, *Pestalotiopsis disseminata*, *Colletotrichum gloeosporioides* and *R Stolonifer* during Bhodia (August - September); *A. niger, C. cladosporioides*, *A. flavus*, *A. fumigatus*, *R. stolonifer*, *Penicillium chrysogenum* and *P. disseminata* during Kotia (October - Novembber) while *P. chrysogenum*, *C. cladosporioides*, *F. oxysporum*, *R. stolonifer*, *A. niger*, *P. disseminata* and *A. flavus* were the dominant aeromycoflora during the Jaruwa generation (December - January) of Muga silkworm (**Fig. 2c**).

**Fig. 2:**
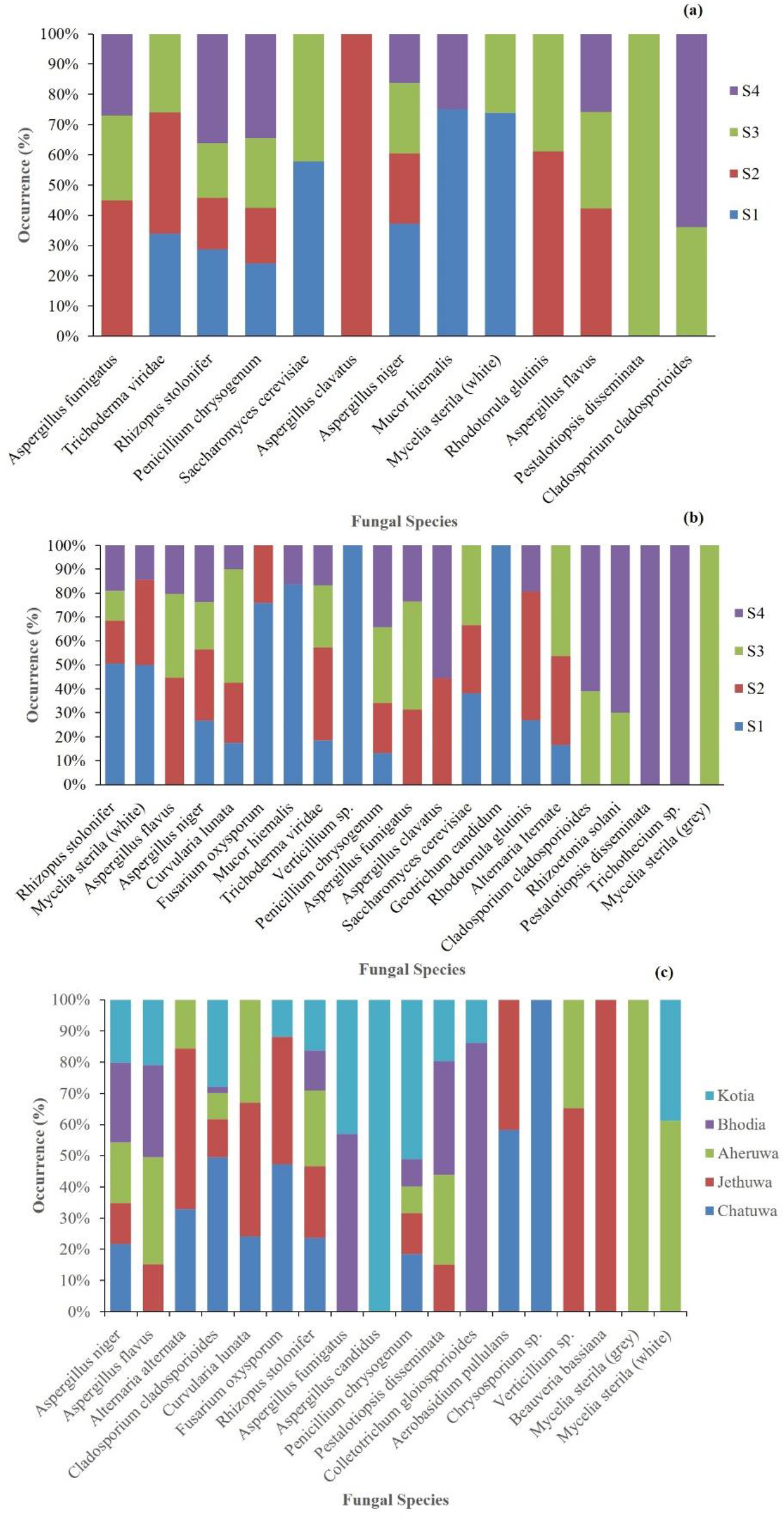
Fungal diversity in (a) non-rhizospheric soil (b) rhizospheric soil and (C) air in the Som plantation area.

#### 3.1.3 Phyllosphere

During Chatuwa generation, *R. Stolonifer* was the dominant fungal species which was found in both the dorsal (23.75%) and ventral surface (32%) of the Som tender leaves (**Fig. 3a**). In semi-mature leaves, *R. stolonifer* was the dominant fungal species (Dorsal: 25.25% and Ventral: 36.5%) (**Fig. 3b**). On both dorsal surface and ventral surface of the mature leaves, *R. stolonifer* was the dominant fungal species encountered (Dorsal: 32.5% and Ventral: 34.5%) (**Fig. 3c**). In Jethuwa generation of Muga silkworm, on dorsal surface of tender leaves *Alternaria alternata* (23.75%) was the dominant fungal species while *Aspergillus flavus* (22.5%) was the dominant fungi on ventral surface of the tender leaves (**Fig. 3a**). In semi-mature leaves, on both dorsal (27%) and ventral surface (22.75%), *R. stolonifer* was the dominant fungal species. In mature leaves *R. stolonifer* (20.5%) was the dominant fungi in the dorsal surface of the leaf while in ventral surface, *Alternaria alternata* (25.25) was the dominant mycoflora. In Aheruwa generation of Muga silkworm, on the dorsal surface of tender leaves *Aspergillus niger* (27.75%) and on the ventral surface *R. Stolonifer* (31%) were the dominant fungi (**Fig. 3a**). In semimature leaves, on both the dorsal and ventral surfaces the dominant fungal species was *R. Stolonifer* (Dorsal: 27.25% and Ventral: 34.25%) (**Fig. 3b**). In mature leaves, *R. stolonifer* were dominant fungal species for both dorsal (32.75%) and ventral surface (28.75%). In *Bhodia* generation, on tender leaves *Aspergillus flavus* was dominant on both dorsal (22%) and ventral surface (25.75%) (**Fig. 3a**). In semi-mature leaves, *A. flavus* (24%) was the dominant mycoflora of dorsal surface while on the ventral surface *A. niger* (18.25%) was the dominant species (**Fig. 3b**). In mature leaves *Pestalotiopsis disseminata* (22%) were dominant on the dorsal surface and *A*. niger(19%) were dominant on the ventral surface of leaves (**Fig. 3c**). In Kotia generation, on tender leaves *A. niger* (D: 29.5% and V: 6.5%) was the dominant fungal species for both the surfaces (**Fig. 3a**). In semi-mature leaves *A. niger* was the dominant fungal species on both the dorsal (27.75%) and ventral surfaces (29.75%) of the leaves (**Fig. 3b**). In mature leaves, *A. niger* was dominant fungal species for both the dorsal (28.5%) and ventral surfaces (30.5%) (**Fig. 3c**). In Jaruwa generation *R. stolonifer* was dominant fungi for both the dorsal (24.75%) and ventral surfaces (31.5%). In semi-mature leaves *R. stolonifer* was the dominant fungi for both the surfaces (D: 39.25% and V: 33%) (**Fig. 3b**). In mature leaves dorsal surface, *R. stolonifer* was the dominant fungal species for both the dorsal (20.25%) and ventral surfaces (23.5%) of the leaves (**Fig. 3c**).

**Fig. 3:**
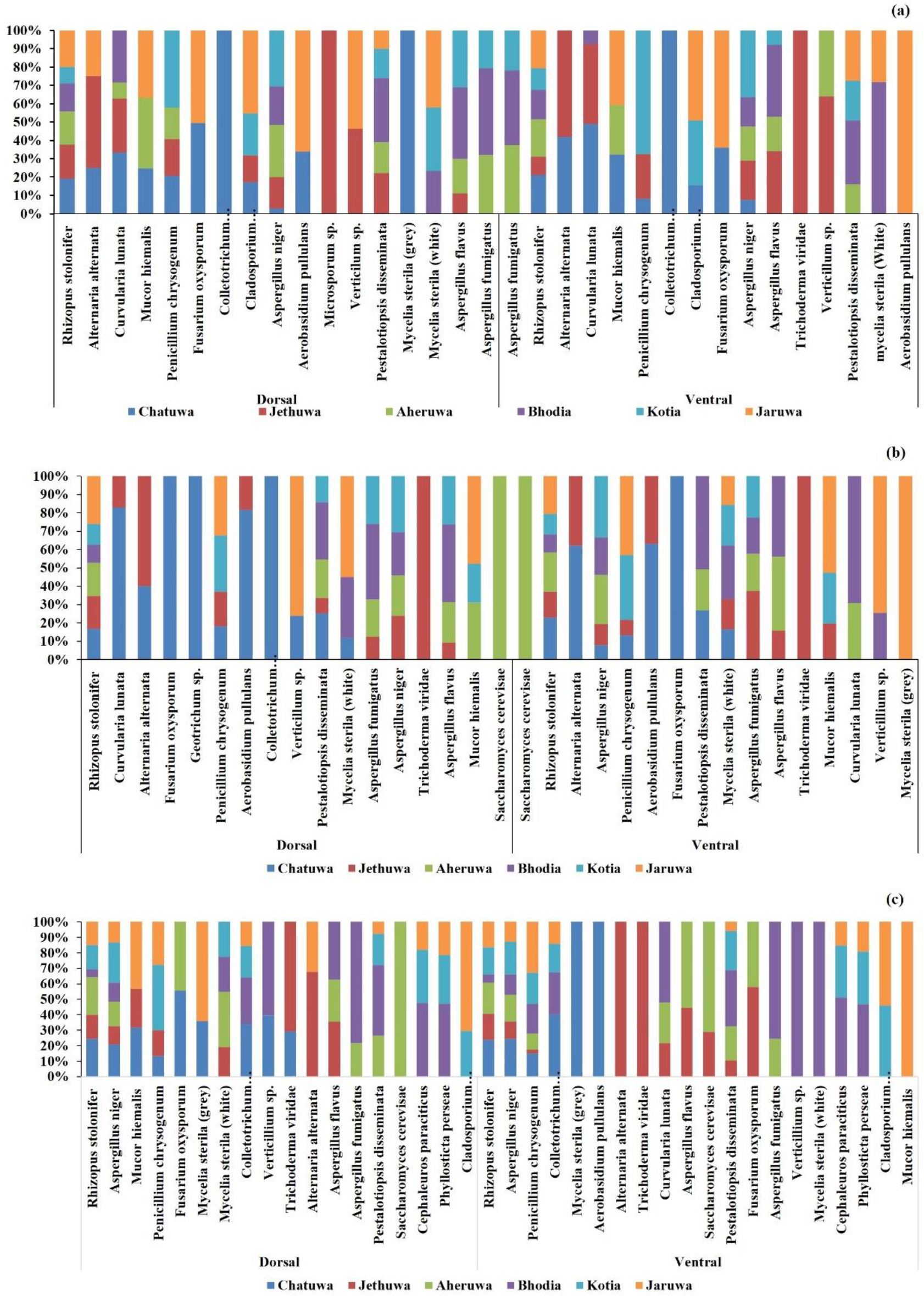
Fungal diversity of phyllosphere in six different generation of (a) Tender, (b) Semi-mature and (c) mature leaves of Som plant.

### 3.2 Comparative composition of the fungal population

#### 3.2.1 Common population

A comparative community analysis of a fungal population of the four environments *viz*. rhizosphere, non-rhizosphere, air and phylloplane using Venn diagram showed 3 fungal species (10%) *viz. Geotrichum candidum*; *Rhizoctonia solani*; *Trichothecium sp*. are unique and exclusive to the only rhizosphere, while there is no fungal species is exclusive to the non-rhizospheric environment. *Aspergillus clavatus*; *Rhodotorula glutinis* are two (6.7%) common species among the rhizosphere and non-rhizosphere. There were no exclusive common species among the rhizosphere and phylloplane while three species (10%) were found common among the rhizosphere, non-rhizosphere and phylloplane *viz. Trichoderma viridae*; *Saccharomyces cerevisae* and *Mucor hiemalis*. Likewise, there were no exclusive species common to non-rhizosphere and phylloplane. Four fungal isolates (13.3%) viz. *Microsporum sp*.; *Geotrichum sp.; Cephaleuros parasiticus* and *Phyllosticta perseae* were unique to phylloplane. There were five common species between the Rhizosphere, Phylloplane and Air *viz*. *Fusarium oxysporum*, *Curvularia lunata*,*Verticillium sp*., *Alternaria alternata* and *Mycelia sterila (Grey)*. Among all the environments, there were eight common species (26.7%) in “Rhizosphere”, “Non-Rhizosphere”, “Phylloplane” and “Air” *viz. Aspergillus fumigatus, Rhizopus stolonifer, Penicillium chrysogenum, Aspergillus niger*, *Mycelia sterila* (white), *Aspergillus flavus*, *Pestalotiopsis disseminata, Cladosporium cladosporioides*. There was no exclusive species between Non Rhizosphere, Phylloplane and Air. Two fungal species were exclusively common among Phylloplane and Air *viz. Colletotrichum gloeosporioides*, *Aerobasidium pullulans*. Three fungal isolates (10%) were exclusively found in the Air *viz. Aspergillus candidus*, *Chrysosporium sp*., *Beauveria bassiana* (**Fig. 4**).

**Fig. 4:**
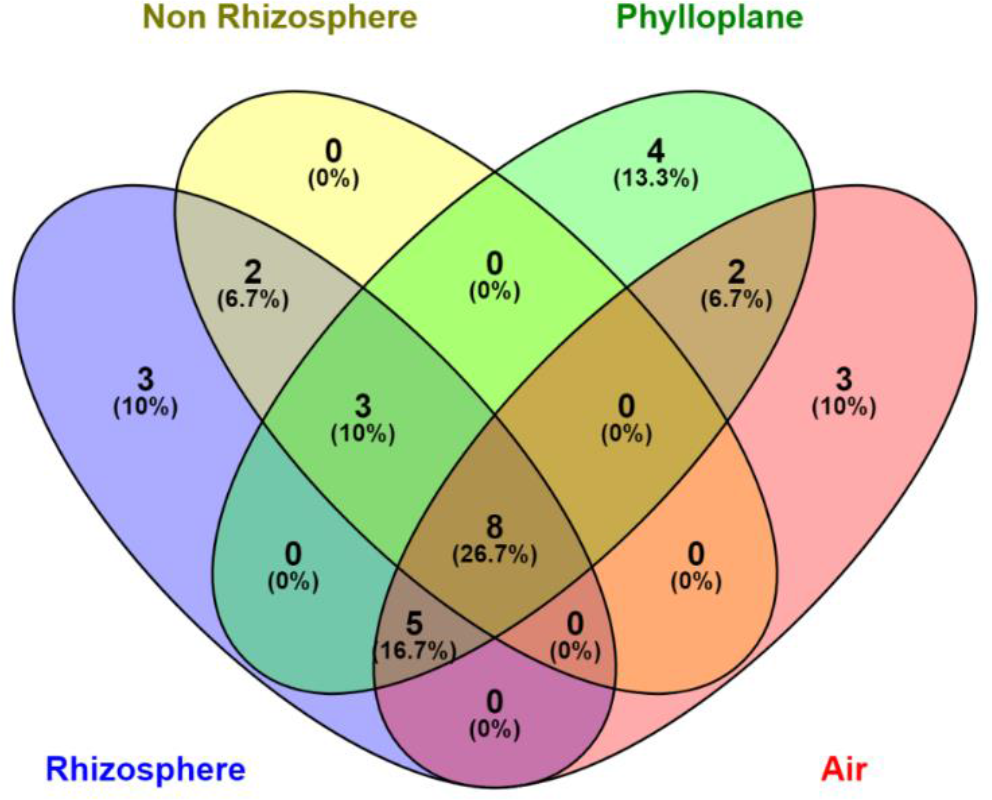
Common population among the four environments; rhizosphere, non-rhizosphere, air and phylloplane.

#### 3.2.2 Comparative population

Comparative population analysis using heatmap showed the rhizosphere and aerosphere were most rich in fungal diversity as compared to non-rhizosphere and phylloplane. The principal component analysis showed the fungal diversity in the rhizosphere, non-rhizosphere and phylloplane are correlative while diversity in the air is different from the same.

**Fig. 5:**
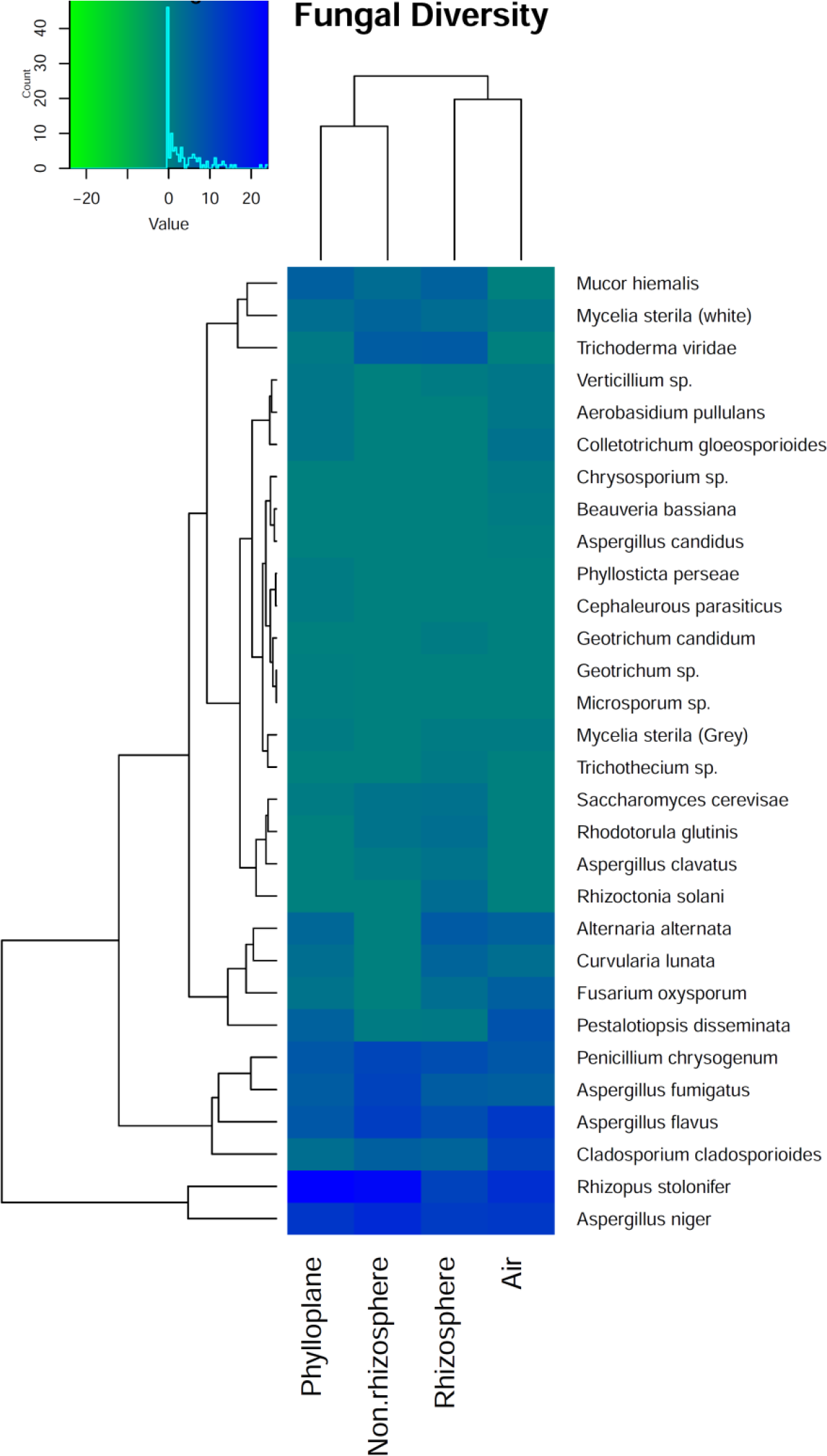
Analysis of compositional difference (a) Heat map constructed with Brey-Curtis dissimilarity index for the fungal diversity of rhizosphere, non-rhizosphere, air and phylloplane, (b) Principal component analysis of the rhizospheric, non-rhizospheric, air and phylloplane mycoflora.

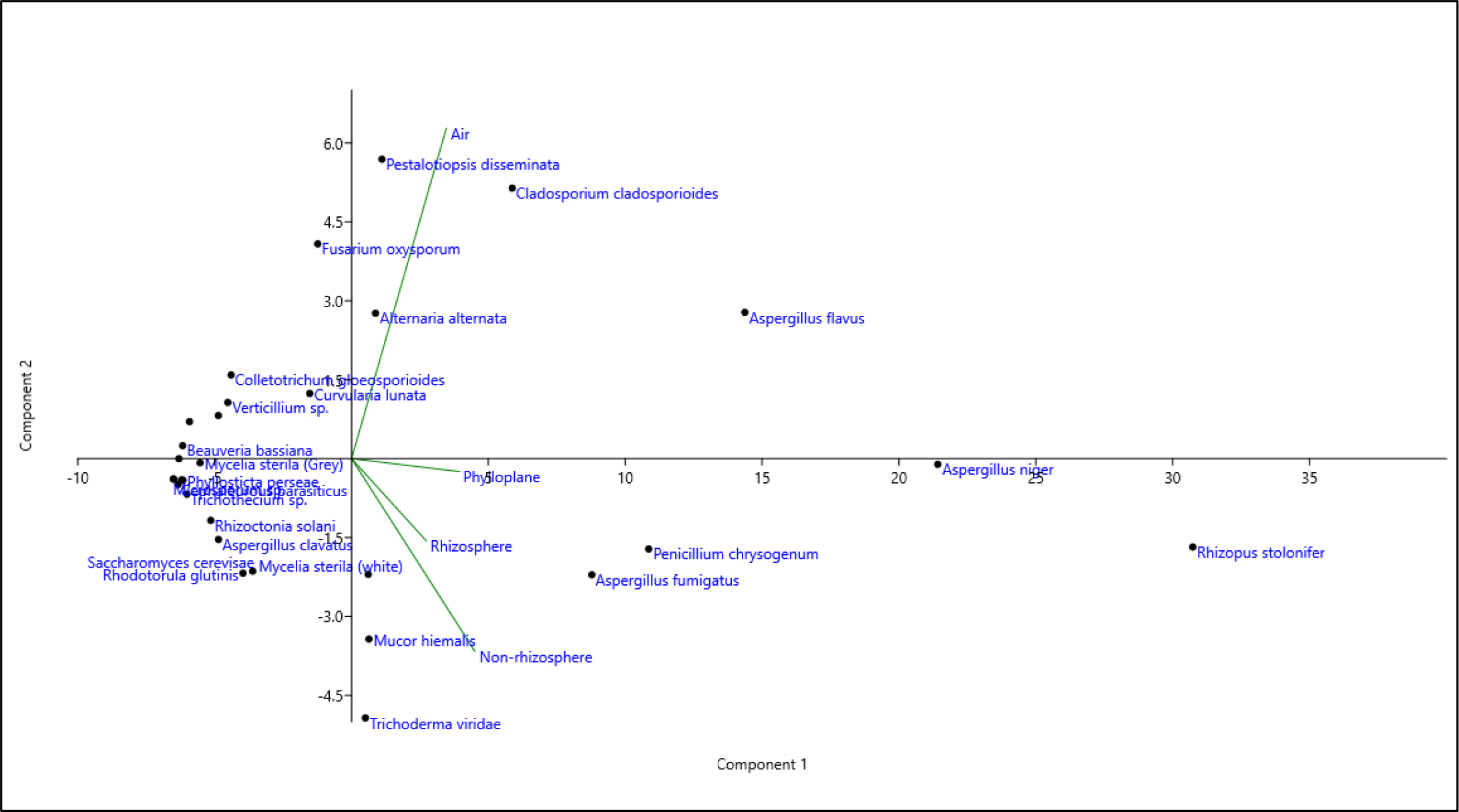

### 3.3 Effect of Environmental Factors on Fungal Diversity

#### 3.3.1 Phylloplane

Canonical correspondence analysis (CCA) between the environmental factors *viz*. Temperature, Humidity, rainfall and total rainy days (**Table. 1**) with the fungal occurrence along with the 6 Muga crop seasons showed that fungal species like *Rhizopus stolonifer*, *Cladosporium cladosporioides*, *Fusarium oxysporum*, *Verticillium sp*., *Mucor hiemalis*, *Aerobasidium pullulans and Mycelia sterila(grey)* showed their occurrence during the Jaruwa (December-January), Chatuwa (February-March) and Jethuwa (April-May) generations of Muga silkworm. The climatic factors did not influence the occurrence of these fungal population. While the occurrence of *Aspergillus niger*, *Aspergillus flavus*, *Aspergillus fumigatus*, *Curvularia lunata*, *Pestalotiopsis disseminata*, *Mycelia sterila (white)* and *Cephaleuros parasiticus* were predominant during the Aheruwa (June-July), Bhodia (August-September) and Kotia (October-November) generations of Muga silkworm. The climatic factors such as temperature, humidity, rainfall influences the occurrence of these mycoflora. While for few other fungal species like *Trichoderma viridae*, *Microsporum sp*., *Alternaria alternata* and *Saccharomyces cerevisiae* both the generations of Muga silkworm as well as the climatic factors were not so influencer.

**Table 1.** Environmental parameters of phyllosphere

| Season | Temperature (°C) | Humidity (%) | Rainfall(mm) | Total rainy days |
| --- | --- | --- | --- | --- |
| Chatua<br>(Feb-Mar) | 31 | 84.75 | 208.25 | 2.5 |
| Jethuwa<br>(Apr-May) | 34.25 | 88 | 1351 | 12.5 |
| Aheruwa<br>(Jun-Jul) | 34 | 92 | 4286.25 | 17.75 |
| Bhodia<br>(Aug-Sep) | 36.5 | 92 | 2645.75 | 15 |
| Kotia<br>(Oct-Nov) | 32 | 88.75 | 112.5 | 1.75 |
| Jaruwa<br>(Dec-Jan) | 25 | 81.75 | 112.5 | 1.25 |

- Data are average of all the five study sites.

#### 3.3.2 Soil

CCA plot shows that certain species like *Aspergillus niger*, *Aspergillus fumigatus*, *Curvularia lunata*, *Rhodotorula glutinis*, *Trichoderma viridae*, *Mycelia sterila* (white) were predominant during the summer season when the P^H^ of the soil is basic and amount of available Phosphorus (**Table. 2**) was higher. Similarly, *Rhizopus stolonifer*, *Aspergillus clavatus*, *Penicillium chrysogenum* were predominant in the soil during the winter season where moisture, organic carbon, and water holding capacity were high.

**Table 2.** Physicochemical parameters of Soil of the plantation areas

| Season | pH | Organic Carbon | N<sub>2</sub> | P<sub>2</sub>O<sub>5</sub> | K<sub>2</sub>O | Water Holding Capacity | Moisture |
| --- | --- | --- | --- | --- | --- | --- | --- |
| Summer | 6.136 | 2.19 | 730.18 | 39.238 | 325.92 | 38.34 | 2.462 |
| Winter | 4.732 | 2.44 | 811.202 | 21.798 | 483.1616 | 46.422 | 8.28 |

- Data are average of all the five study sites.

**Fig. 6:**
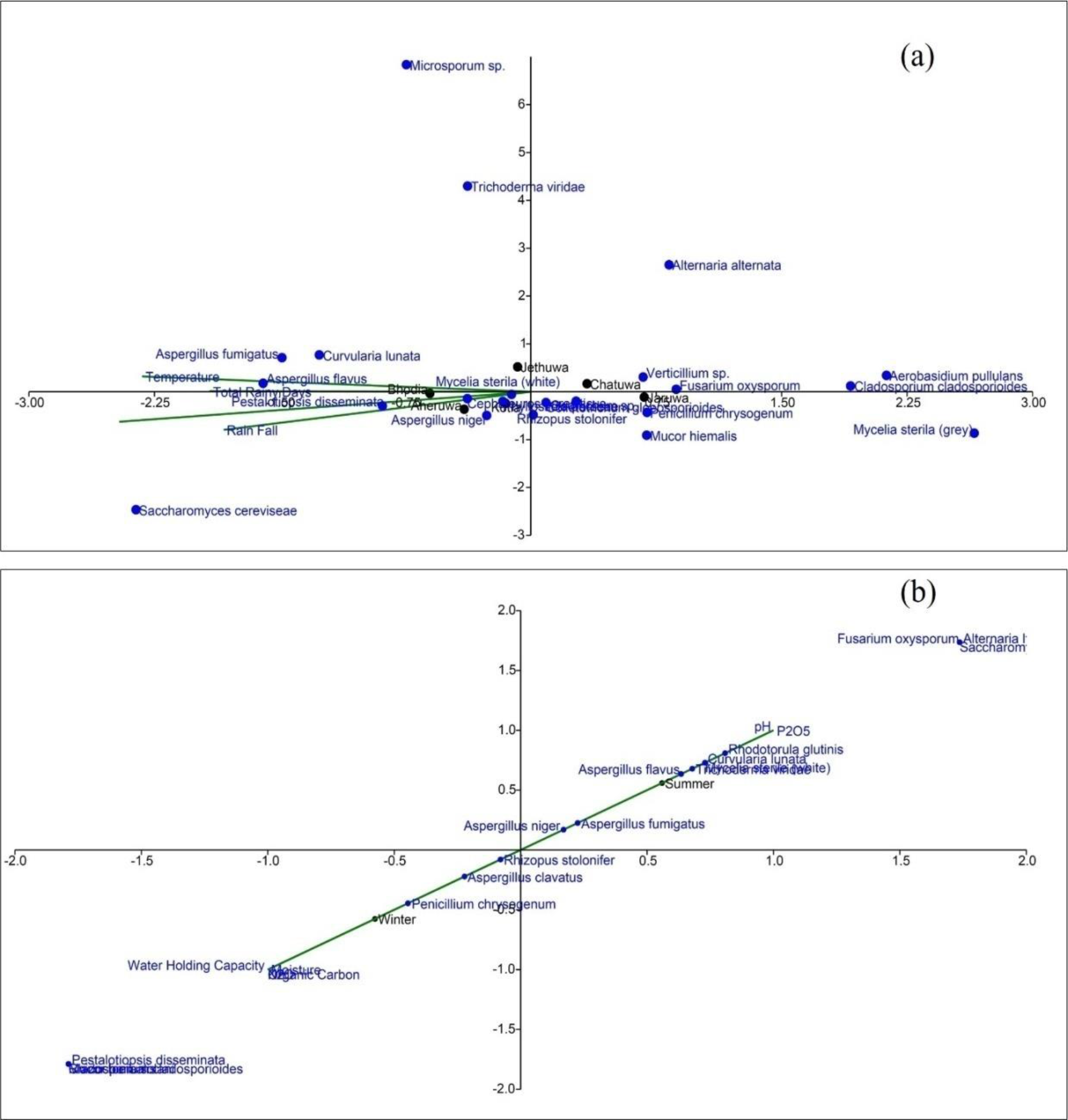
Canonical correspondence analysis of fungal diversity and environmental variables for (a) Phyllosphere: Phyllosphere diversity was highly dependent on the temperature, rainfall, relative humidity of the place (b) Soil: soil diversity differs in summer and winter.

## 4.0 Discussions

Sericulture is one of the vital agro-based industry which comprises of various inter-linked activities from the cultivation of food plants and maintenance of the leaves for quality cocoon production, reeling of cocoon and processing of the cocoon for production of silk fiber. Sericulture along with silk weaving is the part and parcel of the cultural heritage of the people of North-East India. The number of sericulture village in North-East region of India is about 38,000 and approximately 1.9 lakh families of Assam are socio-economically dependent on this industry. Assam is the only state in the country producing all the varieties of silk (Unni et al., 2009). Among the different silkworm varieties *viz. Bombyx mori* (Mulberry silkworm), *Samia ricini* (Eri silkworm), *Antheraea proylei* (Oak tassar silkworm) and *Antheraea assamensis* Helfer. (Muga silkworm); Muga silk worm or *Antheraea assamensis* Helfer. is endemic to North East Region of India producing the golden colored exquisite silk, unavailable in other parts of the world (Amalendu et al., 2013). Assam itself produces about 86.07 percent (136 tonnes) of India’s total Muga silk (158 tonnes in FY 2014-15) and hence it is considered as the main production zone of Muga silk (Central Silk Board, 2015). Though Assam produces a major proportion of Indian Muga silk but maximum Muga silk production can be observed in upper Assam as so, there is a famous proverb in Assamese, “*Namonir sonch ujanir goch*” which means, seed cocoon from lower Brahmaputra valley reared in upper Brahmaputra valley always ensures the successful harvest of cocoons (Sarmah et al., 2010). In upper Brahmaputra valley of Assam, Dhemaji, Dibrugarh, Lakhimpur, and Sibsagar district produces commercial Muga cocoons in bulk quantities. While in Lower Brahmaputra valley Muga sericulture is practiced in Goalpara, Kamrup and Kokrajhar district only. Mostly Muga seed cocoons are produced in the eastern and western part of Goalpara district and south-western part of Kamrup district (Central Silk Board, 2015). Goalpara district has been given geographical identification mark as the climate is suitable for silkworm rearing (Goswami and Bhattacharya, 2013). Hence there is a maximum possibility to boost up the sericulture industries to a large scale. It will also increase rural employment generation possibilities as well as women empowerment in this area.

Host plants play an important role in sericulture industry. Though Muga silkworm is a polyphagous worm, Som plant or the *Persea bombycina* Kost. remain as the primary host plant (Nath et al., 2008). As of till now a number research has concluded the importance of the Som plant in production of Silk of quality. A healthy nutritious disease-free plant can give rise to healthy cocoon which can reel to a good quality of silk (Neog et al., 2015; Sarmah et al., 2010). Lower Assam already have lesser number of sericulture industrial pockets as compared to the upper Assam and recent urbanization in the region has greatly affected the quality of host plant affecting the quality of silk. Different fungal foliar diseases of Som like grey blight, leaf rust, red rust causes 13.8 - 41.6% yearly loss of leaf yield (Bharali, 1969; Das et al., 2003). So, determining the mycofloral community structure of the plantation area has become the utmost importance to sustain industry for our future generation. In this present study, we have studied the comprehensive fungal distribution in the rhizosphere, non-rhizosphere, air and phylloplane in different Muga seasons.

The rhizospheric and non-rhizospheric mycoflora are greatly influenced by the fertility of the soil, organic matter decomposition, diseases of roots and production of antibiotics. The climatic and edaphic factors greatly influence the distribution pattern of microfungi in the soil of perennial plant (Saksena, 1955; Warcup, 1951). Rhizospheric samples were rich in diversity in comparison to the non-rhizosphere, air, and phylloplane. The rhizosphere soil has the capacity to host more numbers of microbial population due to physiological effect of the root system (Mendes et al., 2013). *Rhizopus stolonifer* (22.13%), *Aspergillus niger* (16.13%), *Aspergillus flavus* (12.12%), *Aspergillus fumigatus* (11.12%), *Penicillium chrysogenum* (10.88%) were the dominat fungal flora in rhizopshere. The major fungal species of rhizosphere is correlative with the non-rhizospheric soils *viz. Aspergillus niger* (12.63%), *Aspergillus fumitgatus* (11.25%), *Rhizopus stolonifer* (11.12%), *Penicillium chrysogenum* (9.5%), *Aspergillus flavus* (9.25%). The similarity in the diversity indicates the linear mobilization of the dominant flora from the non-rhizospheric environment to the rhizosphere or towards the active source of energy and other metabolites (Elaine R. Ingham, 2018). Phylloplane diversity showed correlation with the rhizospheric and non-rhizospheric fungal population with the dominance of *R. Stolonifer* (24.03%), *A. niger* (13.73%), *A. flavus* (7.36%), *P. crysogenum* (7.26%), and *A. fumigatus* (6.31%). The results indicate a cyclic relationship between the mycoflora of non-rhizosphere, rhizosphere, and phyllosphere. The same can be observed from the principal component analysis (**Fig. 5b**) where the mycoflora of rhizosphere, non-rhizosphere, and phylloplane were correlated, while the flora of aerosphere was different with the dominance of *Rhizopus stolonifer* (15.09%), *Aspergillus niger* (13.42%), *Aspergillus flavus* (13.42%), *Cladosporium cladosporioides* (11.38%), and *Pestalotiopsis disseminata* (8.17%). The dominance of *Cladosporium cladosporioides* and *Pestalopsis disseminata* in the air were justified by their airborne nature. *Pestalopsis disseminata* is a ubiquitous fungus in outdoor air environment with maximum spore count in summer (Das et al., 2010). A linear relationship between the temperature, rainfall, relative humidity and population of *P. disseminata* can be observed from the CCA plot (**Fig. 6a**). The density of *P. disseminata* increases with increase in temperature, rainfall, and relative humidity, thus it was recorded least in Jaruwa generation (December - January) when the temperature drops below ≥ 25 °C. *Pestalotiopsis disseminata* is also the major pathogen of Som which causes grey blight (**Supplementary Table. S1**), one of the major disease for which a significant annual loss is recorded every year (Das et al., 2003).

Seasonal variation plays an important role in shaping the fungal community structure, as temperature, humidity, rainfall, and concentration of different minerals changes with seasonal change which affect the diversity. In summer, the soil has the highest dominance of *Aspergillus flavus*, *Aspergillus niger*, *Trichoderma viridae*, *Alternaria alternata* while winter season favors the occurence of *Penicillium chrysogenum*, *Cladosporium cladosporioides*, *Aspergillus niger*, *Rhizoctonia solani*, and *Rhizopus stolonifer* (**Fig. 6b**). The phyllosphere of Som also showed variation in fungal diversity in accordance to Muga rearing season, though the principle population remains the same. The *Rhizopus stolonifer* was present in all the generation of Muga except Kotia but it was the major dominant flora of Chatuwa (February - March) and Jaruwa (December - January) generation. It was present in both dorsal and ventral surface of all the three leaf types ie. tender, semi-mature and mature. The Jethuwa generation (April - May) showed the dominance of *Alternaria alternata*, *Aspergillus flavus*, and *R. stolonifer*. This shift in the overall diversity of fungus is due to the gradual rise in the environmental temperature, as the *Alternaria* and *Aspergillus* favour the summer environment (de Ana et al., 2006). Subsequently, the later generations of summer like Aheruwa (June - July), Bhodia (August - September) and Kotia (October - November) were dominated by the fungus *Aspergillus niger* and *Aspergillus flavus* though *R. stolonifer* was ubiquitous to all the generation. Kotia generation (October - November) showed the complete dominance of *Aspergillus niger*. Mature leaves of Bhodia generation showed the presence of *Pestalotiopsis disseminata* on the dorsal surface of the leaf and it is the major pathogen of Som for causing grey blight. The presence of *Pestalotiopsis disseminata* in Bhodia generation is due to the favorable temperature and favorable relative humidity. It was earlier reported that a temperature of 28 ± 5 °C and RH of 70 - 80% favors the dissemination of *P. disseminata* (Das et al., 2010). Few works on Som plant also describes the Bhodia and Aheruwa generation were highly prone to grey blight(CMER&TI, 2007).

## 5.0 Conclusion

This study gives the qualitative and quantitative data on non-rhizospheric & rhizospheric soil, air and phylloplane mycoflora of Som (*Persea bombycina* Kost.) from Goalpara district of Assam. The present study provides a comprehensive report on the diversity of mycofloral during all the six generations of Muga silkworm in a year i.e. the Jaruwa (December-January), Chatuwa (February-March), Jethuwa (April-May), Aheruwa (June-July), Bhodia (August-September) and Kotia (October-November). Diseases are always an output of three factors, host, environment and pathogen. The present study describes how different environmental factor like temperature, humidity and rainfall affects the occurrence and diversity of fungi in soil, air, and phylloplane of Som plant. Interestingly it was found that the diversity of *Aspergillus niger, A. candidus, A. fumigatus* and *Pestalotiopsis disseminata* were more in summer season or when the temperature is high along with rainfall and humidity. These two genera are involved in diseases of Som viz. Aspergillosis and Grey blight respectively. This study also revealed the physicochemical properties and texture of soil samples of Som plantation area of the district. The findings of this study provide a deep insight into the mycofloral of pedosphere, atmosphere, and phyllosphere. This knowledge could be used to decide and chalk out different disease management protocol to support and sustain the silk industry for future generations.

## Acknowledgment

The authors are grateful to DBT-Advanced Institutional Biotech Hub, BN College, Dhubri for providing the basic infrastructure to carry out the research work. Authors are thankful to Department of Biotechnology, Govt. Of India for the financial assistance.

## Authors Contribution

Equal contribution to all the authors.

